# ScTree: Scalable and robust mechanistic integration of epidemiological and genomic data for transmission tree inference

**DOI:** 10.1101/2024.11.20.624621

**Authors:** Hannah Waddel, Katia Koelle, Max S.Y. Lau

## Abstract

Phylodynamic models capture joint epidemiological-evolutionary dynamics during an outbreak, providing a powerful tool to enhance understanding and management of disease transmission. Existing phylodynamic approaches, however, mostly rely on various non-mechanistic or semi-mechanistic approximations of the underlying epidemiological-evolutionary process. Previous work by Lau *et al*. [1] has shown that full Bayesian mechanistic models, without relying on these approximations, can enable highly accurate joint inference of the epidemiological-evolutionary dynamics including the unobserved transmission tree [1, 2]. However, the Lau method faces major computational bottlenecks. As the volume of genomic data collected during outbreaks continues to grow, it is crucial to develop scalable yet accurate phylodynamic methods. Here we propose a new Bayesian phylodynamic model, overcoming the major scalability issue in the Lau 2015 method and enabling a readily deployable, yet accurate, phylodynamic modeling framework. Specifically, we develop a *sc*alable spatio-temporal phylodynamic framework for inferring the transmission *tree* (*ScTree*) and other key epidemiological parameters considering the infinite sites assumption in modeling mutation on the sequence level, in contrast to Lau 2015 in which mutation was modeled explicitly on the nucleotide level. Our approach features full Bayesian implementation utilizing a realistic likelihood to mechanistically integrate epidemiological and evolutionary processes. We develop a computationally-efficient data-augmentation Markov Chain Monte Carlo algorithm, inferring key model parameters and unobserved dynamics including the transmission tree. We assess performance of our method using multiple simulated outbreak datasets. Our results indicate that our method can achieve high inference accuracy, comparable to the performance of the Lau 2015 method. Additionally, our method scales significantly more efficiently for large outbreaks, with computing time increasing linearly with outbreak size, compared to the exponential scaling of the Lau method. We also demonstrate our method’s utility by applying our validated modeling framework to a dataset describing a foot-and-mouth disease outbreak in the UK [3]. Our results show that our method is able to generate estimates of the transmission dynamics consistent with those from the Lau 2015 method, further demonstrating the robustness of our new approach. In summary, our method provides a computationally-efficient, highly scalable, accurate modeling framework for inferring the joint spatio-temporal dynamics of epidemiological and evolutionary processes, facilitating timely and effective outbreak responses in space and time. Our method is implemented in our R package *ScTree*.

**Author summary:** Phylodynamic models integrate epidemiological and evolutionary dynamics to better understand disease transmission during outbreaks. However, many existing models rely on approximations that limit their accuracy and interpretability. Previous work by Lau et al. has shown that full Bayesian mechanistic models, without relying on these approximations, can enable highly accurate joint inference of the epidemiological-evolutionary dynamics including the unobserved transmission tree. Their method, however, faced significant computational challenges, particularly with large datasets. In this study, we present a new Bayesian phylodynamic model, *ScTree*, designed to overcome the scalability issues of the Lau 2015 method. By adopting the infinite-sites assumption for modeling mutations, rather than explicitly modeling nucleotide-level changes, *ScTree* achieves a significant improvement in computational efficiency while retaining accuracy of model inference. Our method, validated through simulations and real outbreak data, provides results comparable to the original Lau model but at a fraction of the computational cost, demonstrating its scalability and practical application for real-time outbreak responses. *ScTree* is implemented as an R package, making it accessible for further research and public health use.

## Introduction

The introduction of next-generation sequencing techniques continues to decrease the cost and difficulty of genetic sequencing, and pathogen sampling and genetic sequencing are now a routine part of outbreak surveillance and response [4, 5]. This has generated an explosion of available pathogen genetic data, which has continued apace in recent years. For example, millions of pathogen genetic sequences are publicly available on platforms such as GISAID, Genbank or NextStrain [6–8], in addition to private or restricted repositories of genetic sequence information. Practitioners using these datasets require methods that scale to the size of available genetic data, while maintaining accuracy and producing actionable information. This creates an environment conducive to the use of phylodynamic methods, which unify the epidemiological process of disease transmission and evolutionary process of the pathogens to sharpen the inference of the transmission dynamics [9–13]. One variable of particular interest to practitioners is the transmission tree (i.e., who-infected-whom), which can facilitate the estimation of key population-level epidemiological parameters such as the reproductive number [14] and fine-grained transmission dynamics including superspreading events [15, 16].

While methods solely built on phylogeny can capture the relationships between genetic samples in an outbreak, a phylogeny calculated from those samples does not directly represent the transmission tree of that outbreak [17, 18]. For instance, a phylogeny alone will not show the direction of disease transmission, and complications arise when ancestor and descendant pathogens are sampled in the same outbreak [9, 19]. While multiple phylodynamic models explicitly incorporating the underlying transmission dynamics, including the transmission tree, have been proposed, they are subject to certain limitations. First, many phylodynamic methods conduct a two-stage inference, in which the phylogenetic tree is estimated first, and the transmission tree is estimated using that fixed phylogenetic tree [4, 19–21]. This two-stage inference can be problematic, as it assumes that the phylogeny does not depend on the transmission tree. While this can expedites inference and performs well when the primary inferential target is not the transmission tree, this is not reflective of the true relationship between the phylogenetic tree and the transmission tree. To address the limitations of two-stage inferential approaches, other methods simplify parts of the data likelihood and conduct inference using an ad-hoc pseudo-likelihood or composite likelihood, rather than a complete likelihood [22, 23]. However, with these approaches, systematic inference and interpretation of certain epidemiological quantities of interest, such as the timing of the transmission tree, are generally challenging.

In this paper, we develop a fully Bayesian mechanistic phylodynamic model that utilizes a realistic likelihood to describe the underlying joint epidemiological-evolutionary process. In particular, we extend a method previously developed by Lau. *et al* [1]. Briefly, the Lau method developed a custom Bayesian data-augmentation MCMC algorithm [24, 25] which is able to efficiently leverage the complete-data likelihood and explore the high-dimensional parameter space of the joint epidemiological-evolutionary model. As such, it can robustly infer the posterior distributions of model quantities including the unobserved transmission tree (i.e., who-infected-whom) and its timing. It was shown to be able to accurately estimate the joint (epidemiological-evolutionary) process, and performed best in estimating the transmission tree in a study comprehensively comparing multiple phylodynamic methods [2]. However, the Lau method faces major computational constraints due to explicitly modeling mutation on the nucleotide level. Specifically, it requires nucleotide-wise imputation of the entire unobserved transmitted sequences, which could result in a extensively large parameter space including unobserved model variables. Though its accuracy makes it an attractive option to use in disease outbreak transmission tree reconstruction, such nucleotide-wise imputation significantly impedes its computational scalability.

The rapid nature of disease outbreaks as they unfold necessitates methods that can be used quickly and accurately. This paper aims to develop a new phylodynamic model scaling the Lau 2015 method to larger outbreaks, without compromising the accuracy of the inference of the transmission dynamics. We accomplish this by incorporating an *infinite-sites assumption* to describe the evolutionary process, with mutations accumulating within an individual following a Poisson process. Rather than imputing mutations at each base pair in a genetic sequence, we model the genetic mutations between sequences through time. In parallel, we develop an efficient data-augmentation MCMC algorithm for this new model, which enables full Bayesian inference of the transmission dynamics leveraging a complete likelihood describing the observations including the observed epidemiological data and sampled genetic data.

Our methodology is first tested using simulated datasets, where we demonstrate that our new model can achieve similar parameter estimation and transmission tree coverage as the Lau 2015 method. Additionally, our results show that our model scales much more efficiently with increasing outbreak size compared to the Lau 2015 method. Furthermore, our methodology effectively accommodates incomplete sampling of infected individuals, maintaining reasonable accuracy in estimating the transmission tree under moderate sampling coverage. To demonstrate the real-life utility of our methodology, we apply our model to a previously analyzed outbreak of Foot-and-Mouth Disease (FMD) in livestock that occurred in the UK in 2001 [1, 3]. Our results suggest that our new method can reconstruct critical estimates similar to those from the Lau 2015 method. Thus, we demonstrate our model’s ability to perform accurate and scalable phylodynamic inference for disease outbreaks.

## Model and Methods

### Stochastic Epidemiological Process

We model the epidemiological transmission process using a general continuous-time spatio-temporal SEIR framework with Susceptible (S), Exposed (E), Infectious (I), and Removed (R) compartments. Let *ξ*_*S*_(*t*), *ξ*_*E*_(*t*), *ξ*_*E*_(*t*), *ξ*_*R*_(*t*) indicate the sets of individuals in each category at time *t* (Table 1). A susceptible individual *j* ∈ *ξ*_*S*_(*t*) is exposed as a new infection with a stochastic rate from a currently infectious individual *i* ∈ *ξ*_*I*_ (*t*) with rate *βK*(*d*_*ij*_, *κ*). The function *K*(*d*_*ij*_, *κ*) denotes a spatial kernel function, which allows the infectious challenge from individual *i* to *j* to vary with distance *d*_*ij*_. For this work, we assume an exponentially-decaying 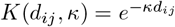, though other kernel options are possible. The overall probability of *j* becoming exposed to the pathogen in the time interval [*t, t* + *dt*) is

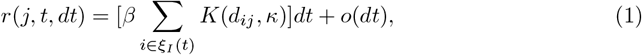

where *o*(*dt*) represents a term that becomes negligible when *dt* is infinitesimally small.

**Table 1.** Model notation for observed and unobserved data.

| Notation | Description |
| --- | --- |
| $\chi_U(t)$ | Indices for the unexposed individuals at time $t$ |
| $\chi_E(t)$ | Indices for individuals exposed by time $t$ |
| $\chi_I(t)$ | Indices for infectious individuals by time $t$ |
| $\chi_R(t)$ | Indices for individuals who have recovered or been removed at time $t$ |
| $\chi_{E \setminus I}(t)$ | Indices of those exposed but not yet infectious by time $t$ |
| $\chi_{I \setminus R}(t)$ | Indices of those infectious but not yet recovered/removed by time $t$ |
| $E_j$ | Time of exposure for individual $j$ |
| $I_j$ | Time of becoming infectious for individual $j$ |
| $R_j$ | Time of recovery or removal for individual $j$ |
| $S_j$ | Time of pathogen genetic sampling for individual $j$ |
| $\psi_j$ | Source of infection for individual $j$ |
| $\delta_{j,m}$ | Genetic distance (Hamming distance) for individual $j$ during interval $m$ between two sampling or transmission events |
| $\Delta_{i,j}$ | Observed genetic distance between individual $i$ and $j$'s genetic samples |

After an individual is exposed, their sojourn time spent in the exposed category (class *E*) is modelled using a *Gamma*(*a, b*) distribution with shape *a* and scale *b*, with a density *f*_*E*_(*a, b*) and cumulative distribution *F*_*E*_(*a, b*) (Table 2) [26, 27]. After this latent period has elapsed, the individual moves into the infectious category (class *I*) and spends an amount of time governed by a *Weibull*(*c, d*) distribution with shape *c*, scale *d*, density *f*_*I*_ (*c, d*) and distribution *F*_*I*_ (*c, d*) [26, 27]. Following this sojourn time in the infectious category, the individual recovers or is removed from the population (class *R*). Note that these sojourn time distributions are selected as appropriate, and do not necessarily need to be the Gamma or Weibull distributions. The sojourn times are assumed to be independent between individuals.

**Table 2.**
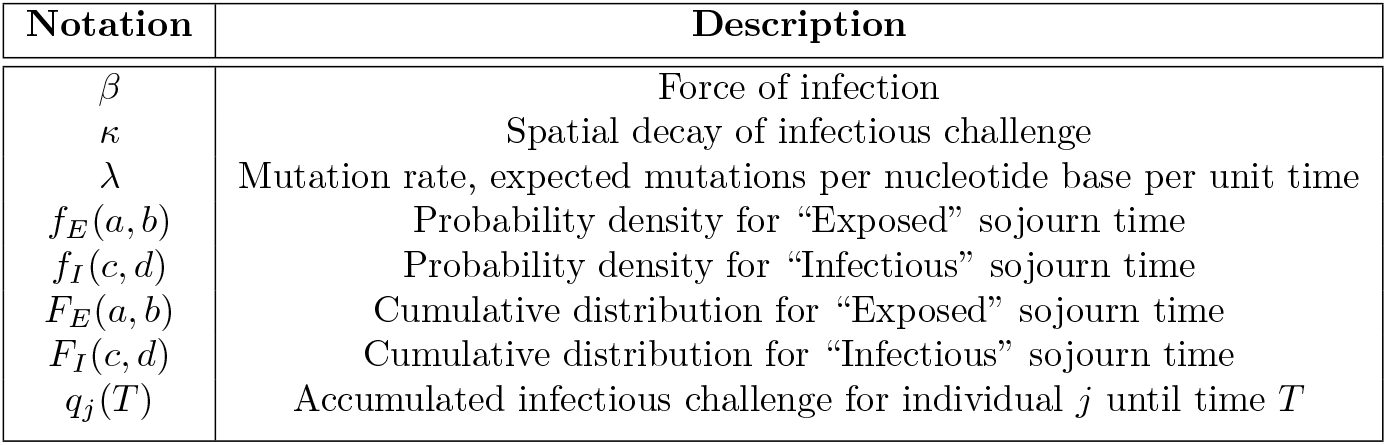
Notation for model parameters.

### Stochastic Evolutionary Process

#### Surrogate Modeling of Evolutionary Dynamics: Infinite-sites Model

Our model aims to mechanistically and fully capture the joint epidemiological-evolutionary process schematically illustrated in Fig 1, which requires inference of unobserved transmission and partially-observed evolutionary dynamics.

**Fig 1.**
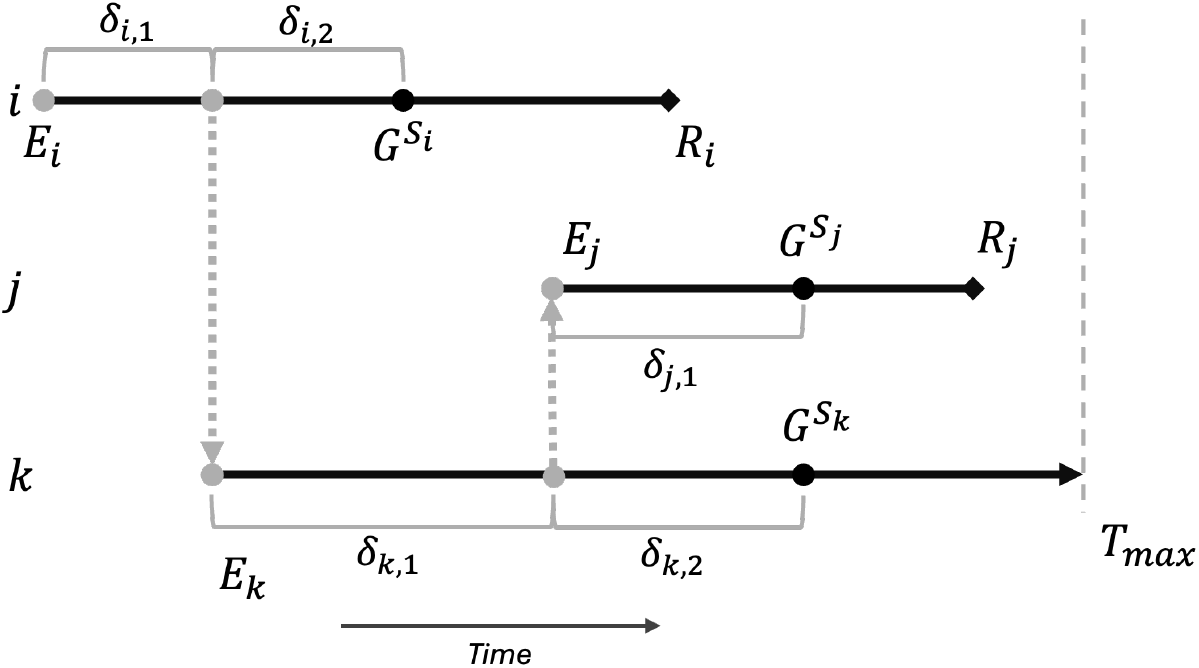
A small outbreak in which individual *i* infects individual *k*, who then infects individual *j*, along with sampled genetic sequences from each individual. Grey circles and lines represent unobserved timepoints and events, such as the exact transmission time from individual *i* → *j*, and the number of mutations that occur between events, denoted by *δ*. Genetic sequences 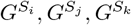 for the individuals are sampled at timepoints *S*_*i*_, *S*_*j*_, *S*_*k*_.

In the Lau method, pathogen evolution is explicitly modeled on the nucleotide level using continuous-time Markov processes such as the Kimura model [28]. Here we describe the evolutionary process using a parsimonious model that adopts the *infinite-sites* assumption, where mutations at each nucleotide site can occur only once, and no mutation reversions can take place. Specifically, we assume that mutations occur according to a Poisson process, where the number of mutations *δ* of a sequence with length *n*, through a period of time *dt* is modeled as *δ* ∼ *Poisson*(*nλ × dt*). The parameter *λ* characterizes the rate of mutations per unit time per site.

It is assumed that this evolutionary process is conditionally independent of the epidemiological process, given the transmission tree and exposure times (modeled by the stochastic epidemiological process previously described).

#### Simulating Ground-truth Evolutionary Dynamics: Nucleotide Substitution Markov Model

Note that the infinite sites assumption offers a surrogate modeling approach for more complicated Markov nucleotide substitution processes [29]. To test the robustness of our surrogate model incorporating the infinite-site assumption, in simulation studies (described in Results) we adopt a two-parameter continuous-time Kimura Markov model [28] used in Lau 2015 as the ‘ground-truth’ model to simulate fine-grained nucleotide-level mutations. Briefly, the Kimura model we use is a nucleotide substitution model which assumes that transition mutations (pyrimidine to pyrimidine or purine to purine) occur at a different, typically higher rate than transversion mutations (pyrimidine to purine or vice versa). The model is governed by two parameters, *µ*_1_ and *µ*_2_, which determine the rates of transition and transversion between base pairs *x* and *y* over an interval length *t* according to probabilities

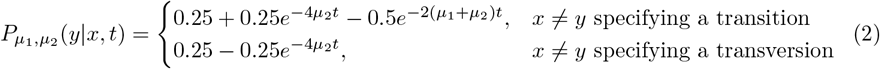

The model allows for reversion of a mutation back into the original nucleotide. Note that, while this model allows substitutions between time points, it assumes no genetic diversity within a host at a given time, following the single-dominant-strain assumption [1].

### A Bayesian Modeling Framework

#### Complete-Data Likelihood

As our inferential procedures make extensive use of *data augmentation* techniques [24, 25], we begin by discussing the formulation of a complete-data likelihood for the joint epidemiological-evolutionary model, assuming all model quantities are known. It is noteworthy that some of the quantities required to calculate the likelihood will be observed directly while others will be inferred/augmented. Notation is explained in Table 1.

We model a population of *N* individuals, for which we observe the geospatial locations of all the individuals. We observe an outbreak in this population between time 0 and time *T*_*max*_. We define *q*_*j*_(*T*) as the accumulated infectious challenge for individual *j* until time *T* :

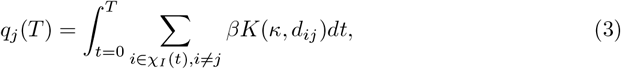

where *K* is the spatial kernel function, and *d*_*ij*_ is the distance between individuals *i* and *j*. The function *P* (*j, ψ*_*j*_) defines the contribution to the likelihood arising from the exposure of *j* by the source *ψ*_*j*_, and is given by

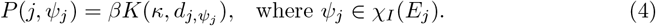

For an individual *j* who is exposed, there is a set of *m*_*j*_ timepoints 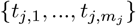, where these timepoints correspond to “critical” events such times of infections and sequence sampling times. Rather than modeling the missing genetic sequences that would be observed at those timepoints, we model the genetic distances between successive sequences. The genetic distance is measured using the Hamming distance, which is the number of base pair differences between sequences [30]. We denote these distances for an individual *j* as *δ*_*j*,._, where *δ*_*j*,._ = {*δ*_*j*,1_, *δ*_*j*,2_, …} is the set of genetic distances between the critical events in individual *j*. (Fig 1).

The contribution to the likelihood from the mutation of an individual *j*’s genetic sequences is given by the following formula:

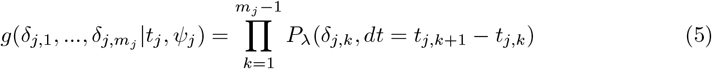

where *P*_*λ*_ is a Poisson distribution with a rate of *nλ × dt*.

Denoting *θ* = (*β, κ, a, b, c, d, λ*) as the scalar model parameter vector, our full likelihood, combining the epidemiological and genomic contributions, is thus given by:

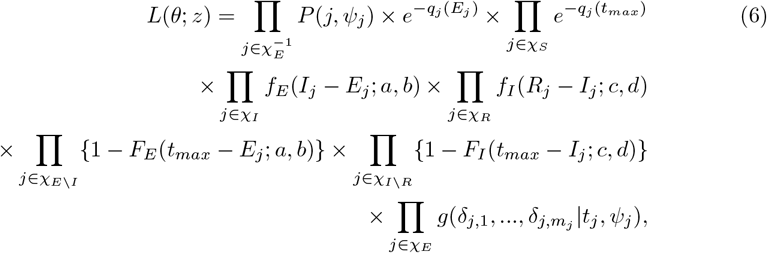

where parameters are defined previously (Tables 1, 2). Note that 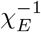 denotes the set of exposed individuals, excluding the earliest exposure or index case.

#### Custom MCMC Inferential Algorithm: Joint sampling of transmission tree, exposure time, and genetic mutations

One major challenge in conducting full Bayesian inference for our previously described joint epidemiological-evolutionary model is developing an efficient MCMC algorithm to explore the vast latent/unobserved parameter space, particularly the joint space of the transmission tree, exposure times, and genetic mutations. Here, we develop a custom MCMC algorithm which can efficiently and effectively explore the vast model parameter space.

We begin by proposing a new infecting source for 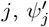, drawn with probability

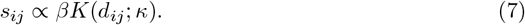

Given the new source 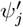, a new exposure time 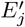 is proposed from a uniform distribution bounded by the infectious times of the infection source and recipient, as well as the genomic sampling time, as an individual would not have a genetic sample before being exposed:

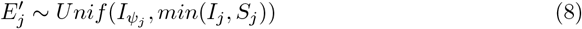

Given the sampled source of exposure 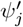 and exposure time 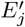, we now describe how we sample the genetic mutations. The main idea is to exploit a *local greedy sampler* that respects and imposes the infinite-sites assumption in the vicinity of the newly proposed exposure time 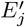. The algorithm is greedy in the sense that it does not necessarily respect the infinite-sites assumption when considering aggregate mutations across all the time points within a particular host. Specifically, our greedy sampler proposes new genetic distances along the *lineage* connecting the observed genetic samples 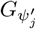 and *G*_*j*_ associated with the newly proposed source 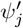 and the individual *j* respectively. Under the infinite-sites assumption along a lineage, a particular *k*^*th*^ genetic distance *δ*_*k*_ neighboring 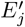 follows a *Binomial* distribution, i.e.,

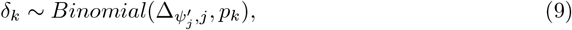

where 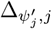 is the genetic distance between the observed samples of 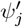 and *j*, and *p*_*k*_ is generally the time duration associated with *δ*_*k*_ normalized by the total elapsed time along the lineage. Fig 2 illustrates our greedy sampler for two typical scenarios. Other scenarios can be readily accommodated and are described in *Supplementary Information*. Note that the acceptance probability for a particular proposed value needs to be properly specified (see *Supplementary Information*).

**Fig 2.**
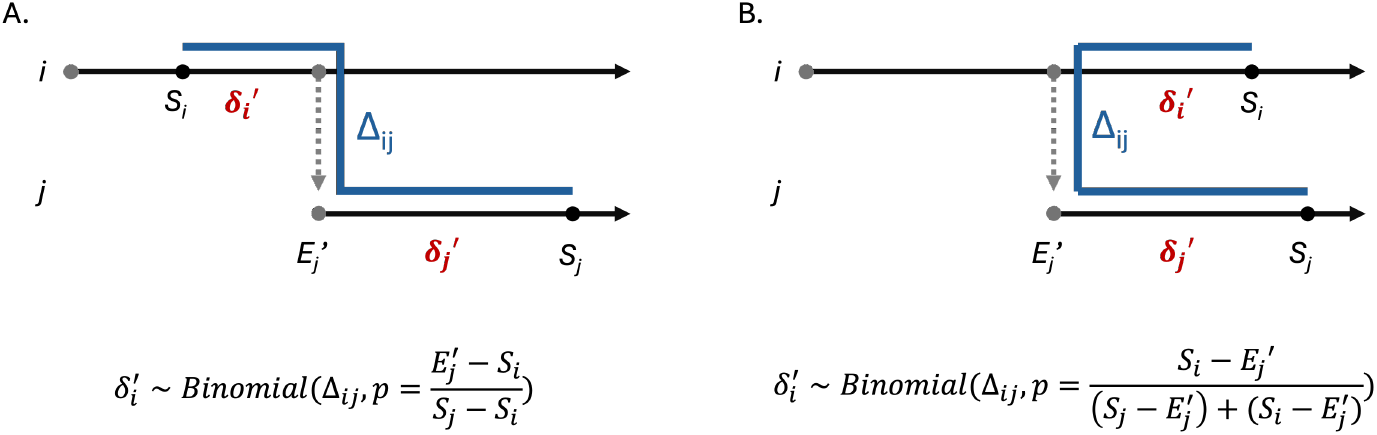
Proposing sequence genetic distances with a locally greedy algorithm. Our local greedy algorithm respects the infinite-sites assumption for mutations neighboring the newly proposed exposure time 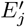, along a lineage (represented by a blue line). We illustrate our algorithm using two scenarios in which both the newly proposed source 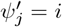 and the infectee individual *j* have observed genetic samples neighboring the newly proposed exposure time 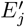. In Scenario A, the genetic sample of 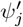 is an ancestor of that of the individual *j*. In Scenario B, the two genetic samples share the same most recent common ancestor.

Further details of our algorithm are described in *Supplementary Information*, including the sampling of the scalar parameters in *θ* = (*β, κ, a, b, c, d, λ*) and the prior distributions adopted.

## Results

### Simulation Studies

We tested our proposed method using multiple simulated synthetic outbreaks. We simulated our outbreaks under the more fine-grained evolutionary model described in Lau 2015, in which a 2-parameter Kimura model [28] is used to model genetic mutation on the nucleotide level (see section *Stochastic Evolutionary Process*). The Kimura model also allows for reversion of mutations through time. By testing our our model, using the infinite-sites assumption, against data simulated under the Kimura model, we can rigorously evaluate the robustness of our surrogate modeling approach under more general genetic mutation conditions.

We simulate datasets based on the simulation studies from the Lau 2015 paper [1]. We consider that an outbreak in a population of *N* = 150 individuals begins at time *t* = 0, given an index case, and proceeds for 104 days. We use epidemiological parameters of *β* = 8, *κ* = 0.02, with sojourn times in the exposed category following a *Gamma*(10, 0.5) distribution and sojourn times in the infectious category following a *Weibull*(2, 2). Pathogen genetic sequences with *n* = 8000 bases are simulated using the mutation parameters of the Kimura model, *µ*_1_ = 5 *×* 10^−4^ and *µ*_2_ = 1.25 *×* 10^−4^. The Lau 2015 model estimates the Kimura model parameters *µ*_1_ and *µ*_2_, while ScTree estimates *λ*, the rate of mutations per day per nucleotide site.

We compare performance by assessing the coverage accuracy of the imputed transmission tree. As in [2], we define coverage accuracy as the proportion of individuals for whom the source with the most posterior support was the true source. We observe in Fig 3 that our new method achieves a comparable coverage rate when compared against the Lau 2015 method, with coverage rate above 90% for each dataset.

**Fig 3.**
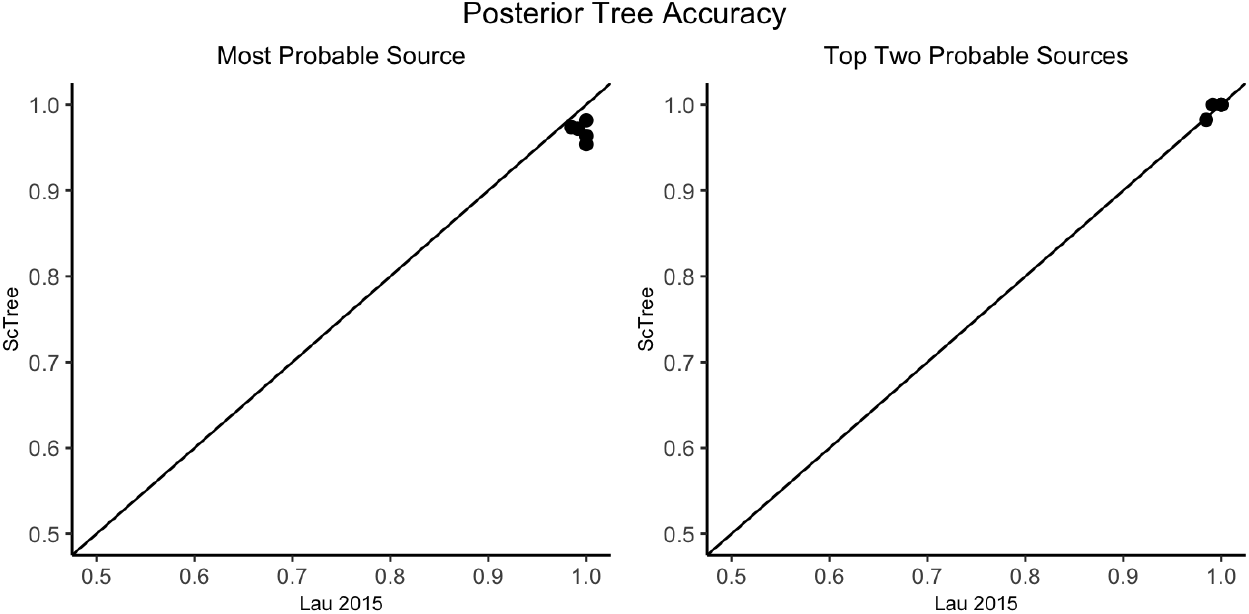
Maximum posterior source coverage accuracy for ScTree versus the Lau 2015 method. The coverage rate here is defined as the proportion of maximum posterior sources which are correct. We see comparable performance by ScTree and the Lau 2015 method. If we expand our source estimates for each method to also include second-most likely sources which are also correct, we see comparable or even improved performance.

We also examined the posterior distributions of scalar parameters for each replicate (Fig 4). We also see similar posterior distributions for the scalar parameters in *θ*, though the estimates for the new method are typically slightly wider (Fig 4). Although the Kimura substitution rate parameters *µ*_1_ and *µ*_2_ are not directly comparable to *λ*, we may approximate the corresponding expected mutation rate (i.e., *λ* under the Kimura model) as *µ*_1_ + 2*µ*_2_. The posterior distribution of our mutation parameter *λ* is also broadly consistent with this approximation, suggesting that our algorithm is able to explore and infer the latent model space efficiently.

**Fig 4.**
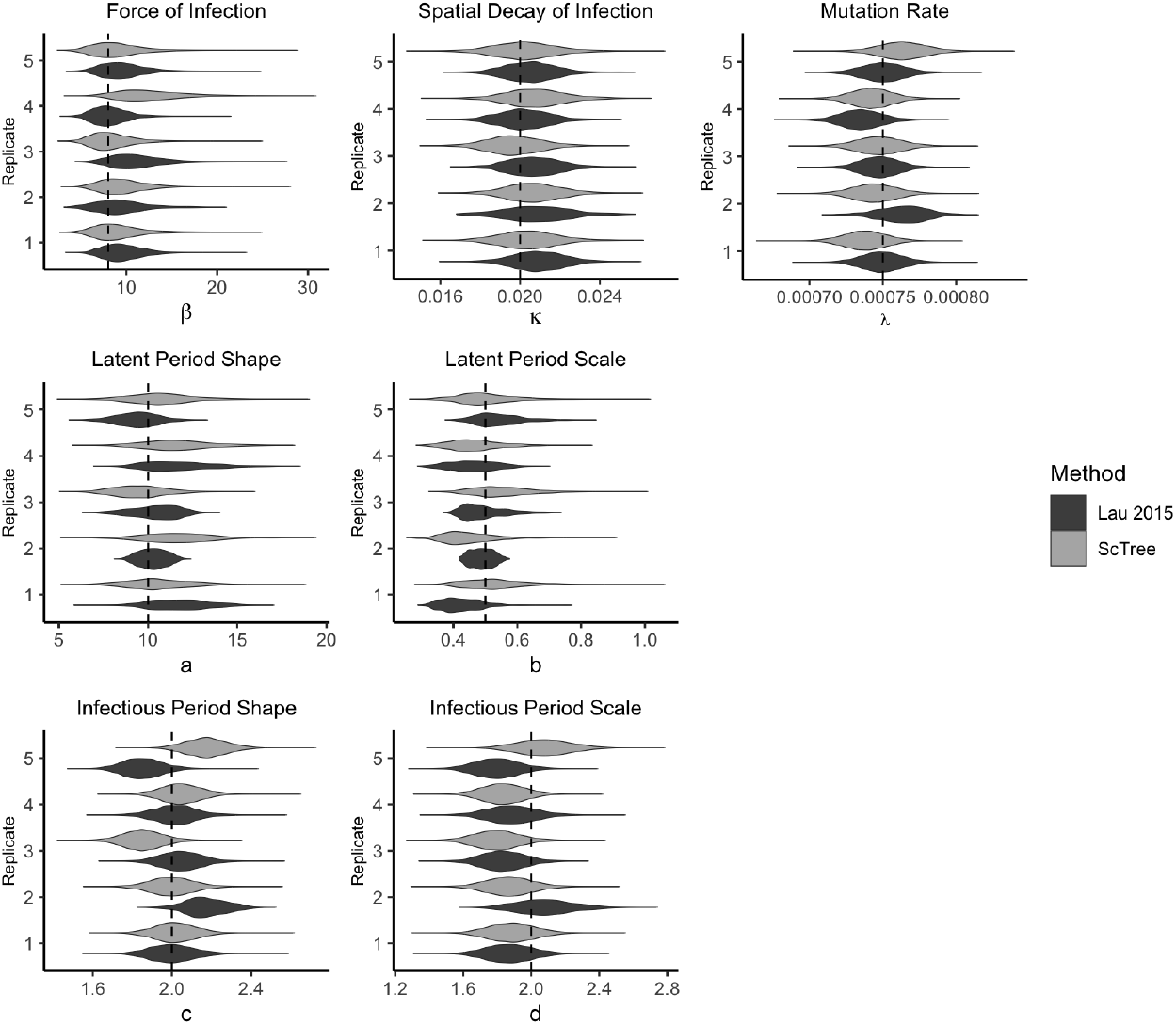
Posterior distributions for scalar epidemiological parameters in the simulation studies. Dashed lines denote the true simulation values of the parameters which are comparable between the Lau 2015 method and *ScTree*. The dashed line shows the approximation value under the Kimura model *λ* ≈ *µ*_1_ + 2*µ*_2_.

### Improved Computational Scalability

We evaluated the computational performance of our method for simulated outbreaks under the same parameter values, with population sizes between *N* = 100 and *N* = 1000 to evaluate computational performance. We simulated a viral genetic sequence *n* = 8, 000 base pairs in length, a similar length to the Foot-and-Mouth Disease Virus and on a similar scale to other viral genome lengths [31, 32].

We see significant improvement in computation time for larger outbreaks when compared to the Lau method, as the computation time for ScTree scales linearly with increasing outbreak size while the computation time for the Lau 2015 method scales non-linearly. For a moderately-sized and densely-sampled outbreak of N=1000 individuals, ScTree computation times are, on average, less than one-fourteenth of the Lau 2015 method (Fig 5). This is expected, as the Lau (2015) method must make 8,000 nucleotide-level imputations for each transmitted sequence, resulting in exploration of the vast joint parameter space of the transmission tree, exposure time, and sequence data. In contrast, our model requires far fewer imputations. We expect the discrepancy in computational performance between our method and the Lau method to further increase beyond the length of the sequence (*n* = 8, 000) being considered here.

**Fig 5.**
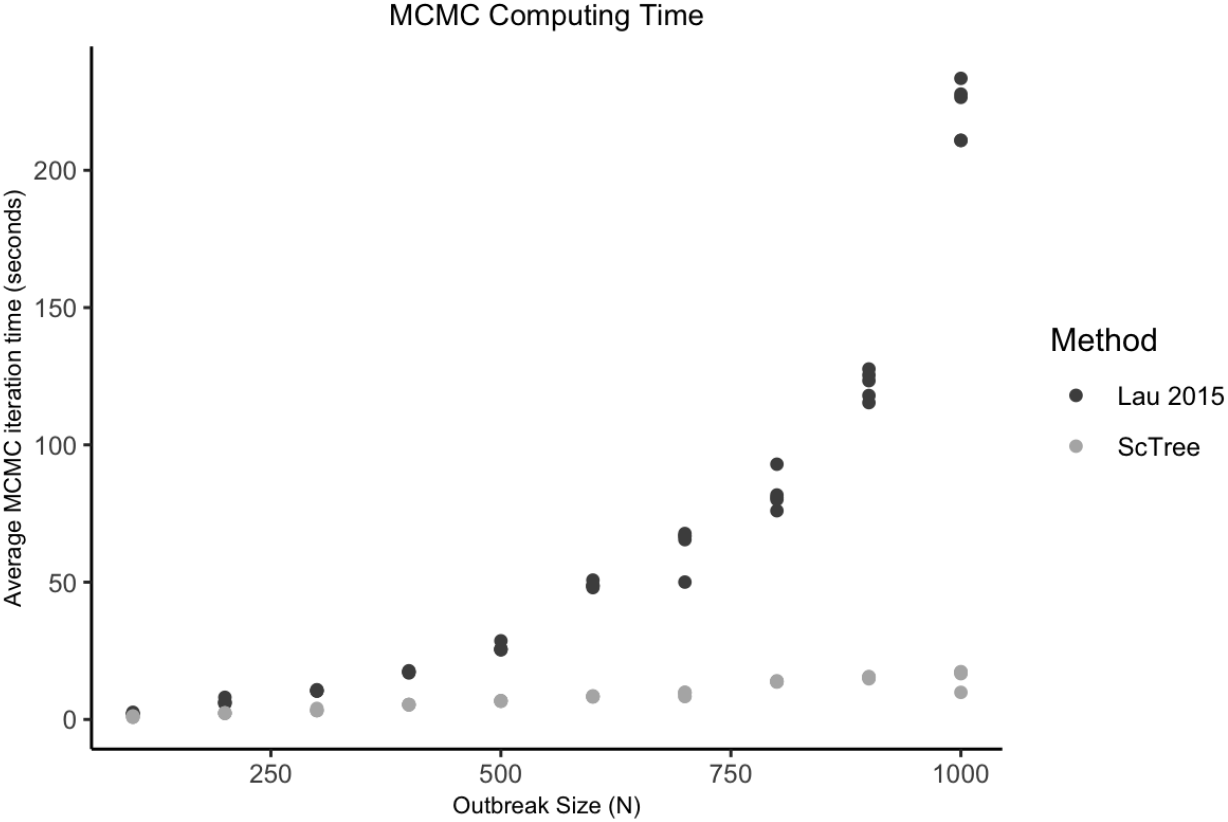
Computation time for 100,000 MCMC iterations for the Lau 2015 method versus ScTree. Among populations of *N* = 100 to *N* = 1000, five outbreaks at each population amount were simulated under the same parameter set.

### Tolerance to Sub-sampling

In real-world conditions, it is possible that only a subset of infected individuals in an outbreak will be sampled. Thus, we tested our method’s tolerance to sub-sampling, where less than 100% of the infected individuals have a genetic sample. Following Lau 2015, we examine the coverage at each MCMC iteration to assess subsampling performance. For each of the five simulated datasets, we considered the baseline scenario with 100% sampling (i.e., every infection has a genetic sample), then subsequently reduced the sampling percentage by randomly removing the genetic samples of some infections and re-ran the inference. We observe in Fig 6 that the tree accuracy of our method decreases as the sampled percentage decreases. This is not unexpected, however, and the coverage of our method still remains around 60%, even as the sampling proportion drops to 10%. This demonstrates that our ScTree model can effectively incorporate available genetic samples to improve inference, while also accommodating a moderate sampling rate in practice and achieving a reasonable estimate.

**Fig 6.**
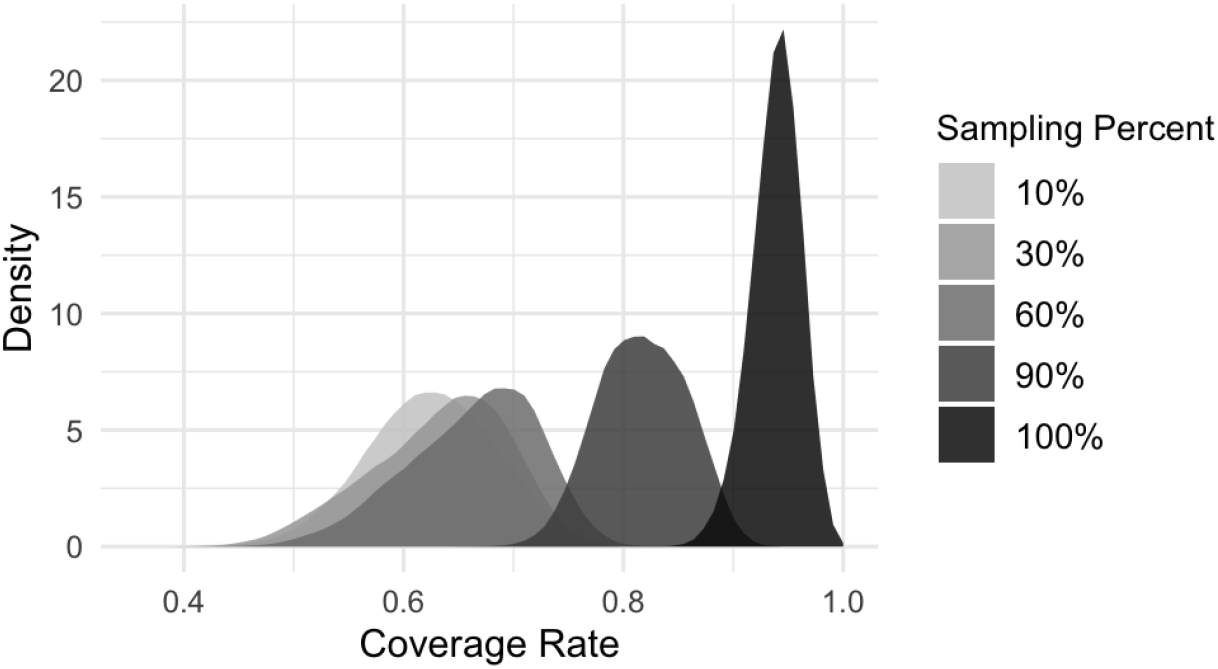
ScTree transmission tree coverage rate under subsampling. Each density plot represents the posterior distribution of the tree coverage rate, pooled from five subsampled datasets derived from the baseline fully sampled datasets, at a specific subsampling rate.

### Case Study: Foot-and-Mouth Disease Virus (FMDV) Outbreak in the UK

We apply our algorithm to a FMDV outbreak which occurred in 2001 in 12 farms in Darlington, Durham County, UK. This dataset was previously analyzed in Lau 2015 and in Morelli et al 2012 [3]. In this case study, following previous work [1, 3], we consider premises with spatially confined host populations as the unit of infection, using the centroids of premises as geographical coordinates. Consensus FMDV genomes sampled from each premise will be used in the inference, with the removal of a premise from the population representing its culling. Each premise was sampled and the outbreak ultimately had 12 genomic samples with a sequence length of *n* = 8196 nucleotides each. The data included geographical location, sampling time and sampled sequences, estimated onset time of lesions (here, taken to be the time of becoming infectious), and removal/culling times of the infected premises.

Fig 7 shows the transmission tree constructed by taking each individual’s most probable posterior source. Our results largely agree with the transmission tree estimated in Lau 2015, and in particular, we reconstruct the longest sequence of transmission (*K* → *F* → *G*→ *I* → *J*) identified in both previous analyses [1, 3].

**Fig 7.**
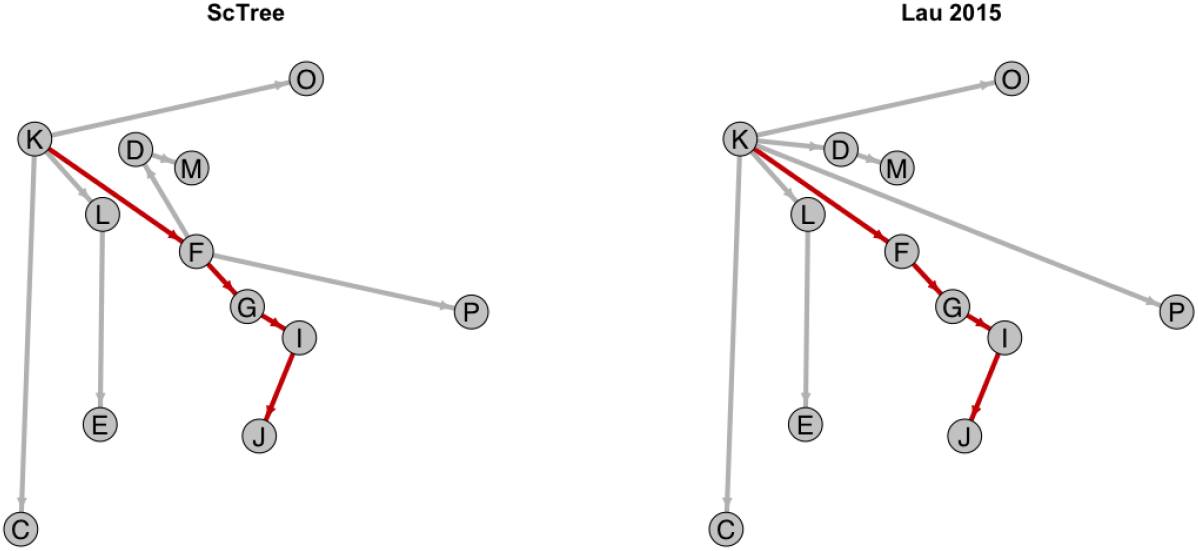
Posterior transmission tree estimates for Foot-and-Mouth Disease outbreak, ScTree (left) versus Lau 2015. (right) The infection source with the highest posterior probability of transmission is used as the estimated source for each farm. The same set of labels for the farms used in [3] is used. We reconstruct the *K* → *F* → *G*→ *I* → *J* transmission chain, the longest sequence of transmissions that was identified both in [3] and [1] (highlighted in red).

Table 3 displays posterior summaries for the model’s scalar epidemiological and mutation parameters. We estimate similar median epidemiological and mutation parameters as the Lau 2015 method, indicating that our method is able to achieve robust estimates of the joint epidemiological-evolutionary dynamics without explicitly considering nucleotide-level mutations. Using the approximation *λ ≈ µ*_1_ + 2*µ*_2_, with the Lau 2015 method we obtained a median estimate of the mutation rate under the Kimura model of 5.98 × 10^−5^ and a 95% credible interval of (4.84 × 10^−5^, 7.27 × 10^−5^). This is broadly consistent with our estimate of *λ* using ScTree, with a median of 7.12 × 10^−5^ and a 95% credible interval of (5.30 × 10^−5^, 9.40 × 10^−5^) (Table 3). We observe that the posterior credible intervals for scalar parameters tend to be slightly wider for our results than for the Lau 2015 method.

**Table 3.** Comparison of the model parameter estimates from the analyses of FMDV outbreak data.

| Parameter | Method | Median (95% CrI) |
| --- | --- | --- |
| $\beta$ (Transmissibility) | ScTree | 0.26 (0.08, 0.76) |
|  | Lau 2015 | 0.16 (0.06, 0.45) |
| $\kappa$ (Spatial kernel) | ScTree | 0.15 (0.03, 0.31) |
|  | Lau 2015 | 0.11 (0.01, 0.25) |
| a (Latent period shape) | ScTree | 1.78 (0.78, 3.67) |
|  | Lau 2015 | 1.77 (0.66, 3.72) |
| b (Latent period scale) | ScTree | 2.83 (1.9, 4.72) |
|  | Lau 2015 | 2.81 (1.87, 5.55) |
| c (Infectious period shape) | ScTree | 4.94 (3.54, 6.96) |
|  | Lau 2015 | 4.93 (3.53, 6.93) |
| d (Infectious period scale) | ScTree | 1.95 (1.2, 2.89) |
|  | Lau 2015 | 1.95 (1.19, 2.89) |
| $\lambda$ (Mutation rate)($10^{-5}$) | ScTree | 7.12 (5.30, 9.40) |

## Discussion

Phylodynamic models capture the joint epidemiological and evolutionary dynamics of an outbreak, providing a powerful tool for enhancing our understanding and management of disease transmission. However, existing phylodynamic approaches have several limitations. In particular, many rely on ad-hoc, non-mechanistic, or semi-mechanistic approximations of the underlying epidemiological-evolutionary process. While these approximations have proven robust when the primary focus is on estimating evolutionary dynamics, systematic inference and mechanistic interpretation of the underlying epidemiological dynamics, particularly the transmission tree, are generally challenging with these approximations.

Lau et al. (2015) made the first attempt of fully mechanistically integrating epidemiological and genomic data within a Bayesian data-augmentation framework. Their methodology is able to utilize a genuine complete-data likelihood that more realistically captures the underlying epidemiological-evolutionary process, as opposed to using ad-hoc pseudo likelihood in many approaches. As such, as shown in a study comprehensively comparing multiple phylodynamic methods [2], their methodology can yield the most accurate estimate of the transmission tree. Their method, however, is limited by poor computational scalability as epidemic size increases. As the amount of genetic data available for outbreaks continues to grow, it becomes imperative to develop a phylodynamic model that not only performs well but is also feasible for use with large, modern outbreak datasets.

In this paper, building on the framework developed by Lau et al. (2015), we develop a more efficient and scalable phylodynamic framework for inferring the transmission dynamics including the transmission tree. Our results show that our method retains the inferential accuracy of the underlying dynamics achieved by the original approach, while significantly reducing the computational burden by bypassing the need to explicitly model mutations at the nucleotide level. Our results also suggest our method can reasonably accommodate the scenarios of incomplete genomic sampling of infected individuals relatively effectively without significantly impacting the tree accuracy.

We also demonstrate our method’s utility by applying our validated modeling framework to a dataset describing a FMD disease outbreak in the UK. Our results show that our method is able to generate estimates of the transmission dynamics consistent with those from the Lau 2015 method, further demonstrating the robustness of our new method. In summary, our method provides a computationally-efficient, highly scalable, accurate modeling framework for inferring the joint spatiotemporal dynamics of epidemiological and evolutionary processes, facilitating timely and effective outbreak responses in space and time.

Our study has several limitations, and future work extending our methodology can be considered. An inherent limitation to our method, as compared to the Lau 2015 method, comes as we work with a summary statistic, the Hamming distance, rather than the nucleotide-level model. There is a trade-off between scalability and precision which requires careful thought to balance in practice. The parsimony in our method results in slightly “flatter” posterior distributions for both scalar parameters and the source of infection. Nevertheless, we observe very comparable performance in inferring the underlying epidemiological-evolutionary dynamics between ScTree and Lau 2015 method.

In addition, the local greedy algorithm, while efficient, does not necessarily apply the infinite-sites assumption globally in the transmission tree. Further work could involve imposing the assumption more globally, such as across a single host or even the entire transmission tree. Finally, our method works with a consensus sequence of the host pathogen populations, which may not show much divergence over a very short time period. Further work with this model may incorporate deep-sequencing data or haplotype networks to better capture within-host population dynamics which are present during an outbreak.

## Supporting Information

### S1 Text. Supporting Information

We present the following supplementary information in the S1 Text: 1) Our general MCMC framework for sampling unobserved data and scalar parameters; 2) Additional scenarios for jointly sampling 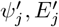, and 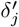 and scenarios in which genetic sampling data is unavailable for a transmission pair; 3) Computing environment.

**S1 Fig. Additional scenarios for sampling sequence genetic distances with a local greedy algorithm when genomic sampling data is available for a transmission pair**

**S2 Fig. Proposing sequence genetic distances with our local greedy algorithm when genomic sampling data is unavailable**.

**S1 Table. Prior distributions for model parameters in simulation analyses**.

**S2 Table. Prior distributions for model parameters in Foot-and-Mouth Disease outbreak analysis**.

## Acknowledgments

We acknowledge the support received through the Emory MP3 (Molecules and Pathogens to Populations and Pandemics) initiative, and Drs. Anice Lowen and David VanInsberghe in the Swine MP3 group for discussions.

## Supporting Information

### SI Text

#### General MCMC framework for sampling unobserved data and scalar parameters

We sample our model quantities including infection source, exposure time, and genetic distances for an individual *j* within a data-augmentation Metropolis-Hastings MCMC framework. The acceptance probability of the jointly-proposed 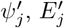, and 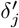, together denoted 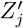 is

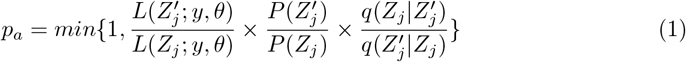

Where 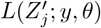 is the likelihood of the unobserved data given the observed data and parameters, P(*Z*_*j*_) is the prior distribution of *Z*_*j*_. The equation 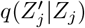 is the proposal distribution of 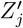 given the current value of *Z*_*j*_.

We update 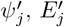, and 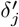 sequentially, and the proposal distributions for 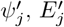, and 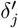 are assumed to be independent, given the other proposed data values. Thus,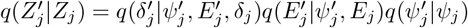. Likewise, for 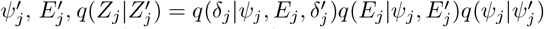, The parameters in *θ* = (*β, κ, a, b, c, d, λ*) are updated sequentially with a random-walk Metropolis-Hastings algorithm. Using *β* as an example, a new value *β*^*′*^ is proposed from a normal distribution centered at the current value of *β* with a variance of 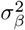.

#### Additional scenarios jointly sampling 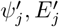 and 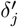

Our main text describes the key details of our MCMC algorithm using two representative scenarios for joint updating of transmission tree, exposure time, and genetic distances. The specific details of the algorithm are determined by availability of sampled genetic data and the timing of the proposed transmission time. Here, we will give further details of our sampling algorithm under other scenarios and when sampling data is unavailable for a transmission pair.

Our proposal scenarios are broadly divided into two categories based on whether there is sample genomic data available for the source-recipient transmission pair. In the main text, we described sampling for 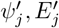 and 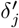 for individual *j* when genomic sampling data is available for the transmission pair and adjacent to the proposed transmission time 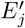. Here, we describe further scenarios which arise, and scenarios in which sampled genomic data is unavailable.

Fig S1 illustrates three scenarios in which genomic data is available for a transmission pair which are not explicitly covered in the main text. We are proposing three genetic distances directly adjacent to the proposed exposure time 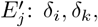 and *δ*_*j*_. In scenario A, for source *i* and infectee *j*, there is a genomic sample for the source taken at time *S*_*i*_, before the infectee’s proposed exposure time 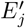. However, in this case, there is a transmission event to some other infectee that occurred between *S*_*i*_ and 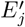 at time *T*_*i*_. To propose the genetic distances *δ*_*i*_, *δ*_*k*_, and *δ*_*j*_ using the sample data, in a local-greedy algorithm, we subtract any “uninvolved” genetic distances which are not adjacent to 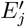 (denoted *δ*_*r*_ in Fig S1 scenario A) to get a “remainder” from the observed sample distance. Then, we sample 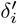 from that remainder using a binomial distribution, while respecting the current genetic distance *δ*_*i*_ into which we are “inserting” 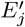 The value of 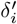 determines the values of 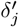 and 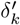. Thus, by this algorithm, we impose the infinite sites assumption locally, in the adjacent genetic distances to 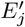. In Fig S1 scenario B, we show the algorithm for proposing genetic distances *δ*_*i*_, *δ*_*k*_, and *δ*_*j*_ when the source genetic sampling time is after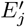. The algorithm is largely the same as in scenario A. In Fig S1 scenario C, we show the case when genetic sampling data is available and we are inserting a new exposure at time 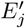 which is the last event for source *i*. The proposal is the same as scenario A, however, we only need to propose two genetic distances.

However, the use of the algorithm described in Fig S1 is contingent upon having a genetic sample for each host, between which we can calculate the genetic distance. We now describe our method for sampling the genetic distances when sample data is not available for the source or the infectee in the transmission pair. Fig S2 shows two sampling scenarios when sample genetic distance data is unavailable. In scenario A, we are inserting a proposed exposure at time 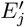 in the middle of source *i*’s events. In this case, we propose the genetic distances adjacent to 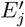 in the source, *δ*_*i*_ and *δ*_*k*_, with a binomial draw and we propose the genetic distance in the infectee, *δ*_*j*_, with a Poisson draw. In Fig S2 scenario B, we are inserting the proposed exposure at the end of source *i*’s events. In this case, we propose the genetic distances adjacent to 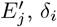 and *δ*_*j*_, with Poisson draws.

#### Computing

We simulate outbreak data (using the Kimura model for the evolutionary dynamics) using the BORIS package, which implements the Lau 2015 model in R. [1]. This work is distributed as an R package ScTree, developed primarily using Rcpp to implement the model inferential algorithm described in our paper [2, 3]. An R package can be found at the following link: https://github.com/hbwddl/ScTree. Data processing, analysis, and visualizations are done via the ape, coda, igraph and ggplot2 packages [4–7].

## SI Figures

**Fig S1.**
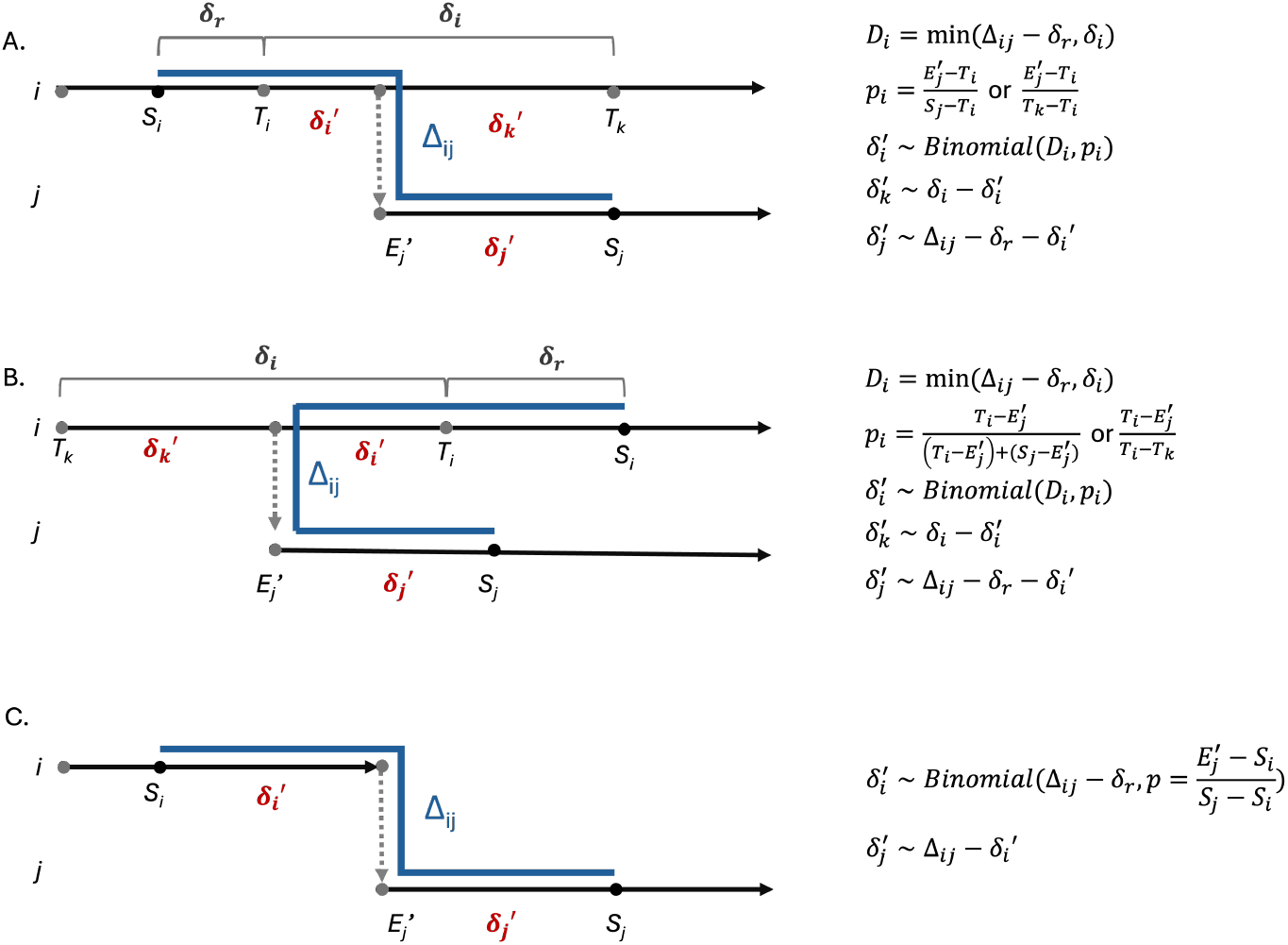
Proposing sequence genetic distances with a local greedy algorithm when genomic sampling data is available for a transmission pair. The local greedy algorithm we use respects the infinite-sites assumption for mutations adjacent to the proposed exposure time 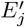. When we have available genomic sampling data, we propose genetic distances adjacent to 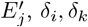, and *δ*_*j*_, with a binomial draw from the sample genetic distance, while respecting the current genetic distances in the source *i*.

**Fig S2.**
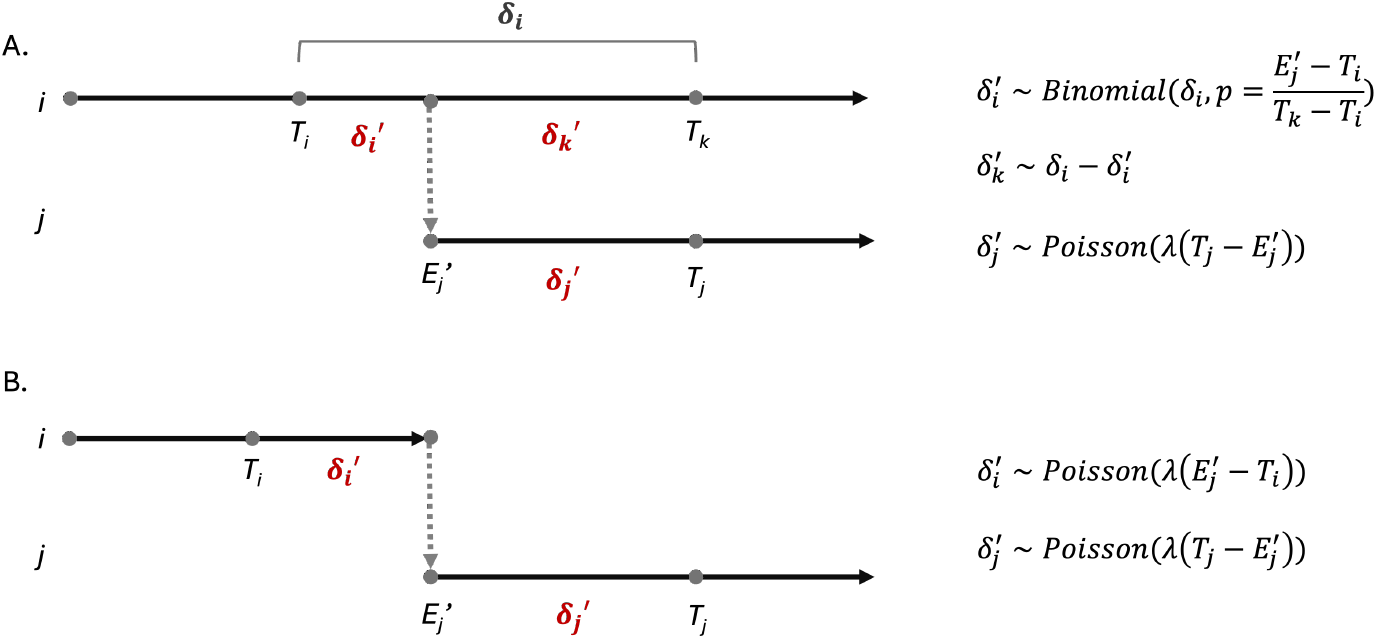
Proposing sequence genetic distances with a local greedy algorithm when genomic sampling data is unavailable. When sample data is not available for the transmission pair *i* and *j*, we propose the new genetic distances adjacent to 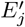 with a binomial draw if we are inserting 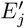 into an existing genetic distance (scenario A), or with a Poisson draw using the current value of *λ* in the MCMC (scenario A, host *j*, and scenario B).

## SI Tables

**Table S1.** Prior distributions for model parameters in simulation analyses.

| Parameter | Method | Distribution |
| --- | --- | --- |
| $\beta$ (Transmissibility) | ScTree<br>Lau 2015 | Unif(0,30) |
| $\kappa$ (Spatial kernel) | ScTree<br>Lau 2015 | Unif(0,10) |
| a (Latent period shape) | ScTree<br>Lau 2015 | Unif(0,50) |
| b (Latent period scale) | ScTree<br>Lau 2015 | Unif(0.1,50) |
| c (Infectious period shape) | ScTree<br>Lau 2015 | Unif(0,100) |
| d (Infectious period scale) | ScTree<br>Lau 2015 | Unif(0,100) |
| $\lambda$ (Mutation rate)($10^{-5}$) | ScTree | Unif(0,50) |
| $\mu_1$ (Transition rate) | Lau 2015 | Unif(0,10) |
| $\mu_2$ (Transversion rate) | Lau 2015 | Unif(0,1) |

**Table S2.**
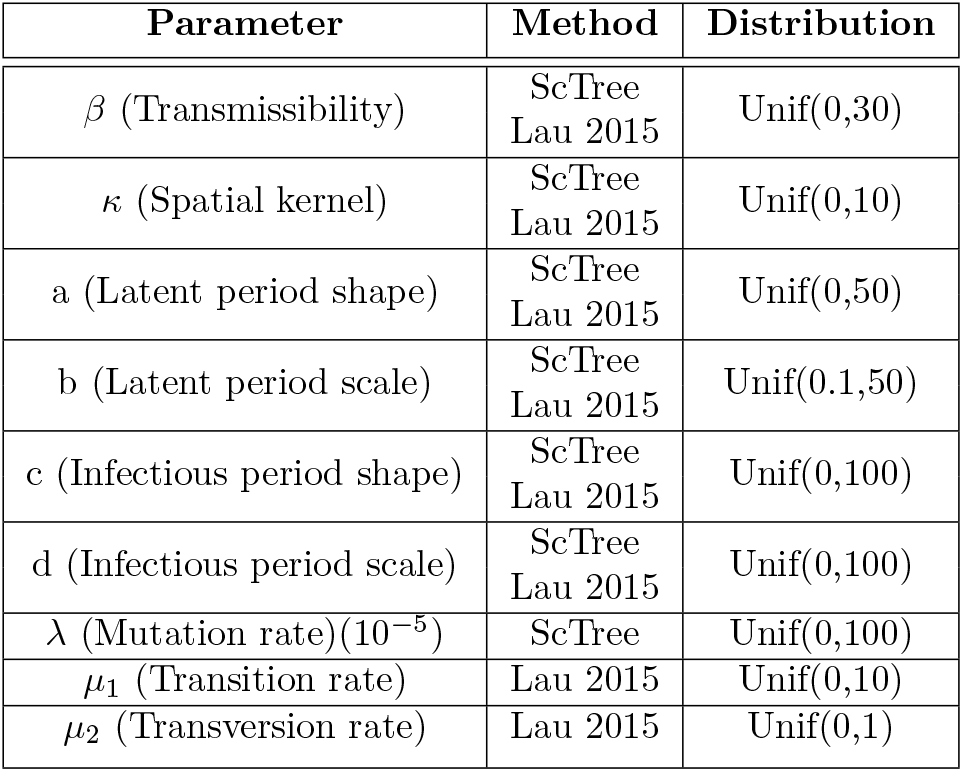
Prior distributions for model parameters in Foot-and-Mouth Disease outbreak analysis.

